# Freewater EstimatoR using iNtErpolated iniTialization (FERNET): Toward Accurate Estimation of Free Water in Peritumoral Region Using Single-Shell Diffusion MRI Data

**DOI:** 10.1101/796615

**Authors:** Abdol Aziz Ould Ismail, Drew Parker, Moises Hernandez-Fernandez, Ronald Wolf, Steven Brem, Simon Alexander, Wes Hodges, Ofer Pasternak, Emmanuel Caruyer, Ragini Verma

## Abstract

Characterization of healthy versus pathological tissue is a key concern when modeling tissue microstructure in the peritumoral area, confounded by the presence of free water (e.g., edema). Most methods that model tissue microstructure are either based on advanced acquisition schemes not readily available in the clinic, or are not designed to address the challenge of edema. This underscores the need for a robust free water elimination (FWE) method that estimates free water in pathological tissue but can be used with clinically prevalent single-shell diffusion tensor imaging data. FWE in single-shell data requires the fitting of a bi-compartment model, which is an ill-posed problem. Its solution requires optimization, which relies on an initialization step. We propose a novel initialization approach for FWE, FERNET, which improves the estimation of free water in edematous and infiltrated peritumoral regions, using single-shell diffusion MRI data. The method has been extensively investigated on simulated data and healthy and brain tumor datasets, demonstrating its applicability on clinically acquired data. Additionally, it has been applied to data from brain tumor patients to demonstrate the improvement in tractography in the peritumoral region.

## 1. INTRODUCTION

Diffusion tensor imaging (DTI), one of the basic frameworks of diffusion MRI (dMRI), is frequently used in clinical studies, because it characterizes tissue microstructure and provides orientation information for delineation of basic fiber tracts using tractography. DTI can be used to enrich radiomic markers of pathology and inform about connectivity in the brain, and is essential for neurosurgical and treatment planning of brain tumors, among others [1]. Since dMRI relies on the measurement of water displacement in tissue, it is confounded by disease processes that cause accumulation of water in the extra-cellular space, such as vasogenic or infiltrative edema. This is further complicated by the presence of contaminants like cancer cells in infiltrated peritumoral regions and parenchymal pathology after brain trauma, any of which can alter the diffusion of water. These processes compromise the specificity of diffusion indices, such as fractional anisotropy (FA) and mean diffusivity (MD), for characterization of white matter and/or tractography, leading to interpretation errors in clinical studies, including those tailored for surgical and treatment planning of brain tumors.

These issues can be alleviated by accurate noninvasive estimation and compartmentalization of water (and other contaminants) in the brain as can be afforded by multi-compartment modeling of the diffusion data. Such modeling captures the uncontaminated tissue as a compartment, and the contaminants as one or more separate compartments. In healthy tissue, multi-compartment modeling is able to alleviate partial volume effects. Multi-compartment modeling allows for more accurate diffusion indices, improved tractography, and novel contrasts such as volume fraction that capture underlying pathophysiological processes [2]. Although the importance of multi-compartment modeling has been recognized [3–7], most of these investigations use multi-shell diffusion acquisitions, since the additional shells allow for more robust and accurate estimation of model parameters. Such acquisitions, however, remain clinically infeasible for the time being.

Single-shell DTI is the predominant type of diffusion MRI acquired in a busy clinical setting. Additionally, large cancer databases, such as the Adult Brain Tumor Consortium (ABTC) (http://www.abtconsortium.org) have single-shell DTI data only. The ability to interrogate the peritumoral manifestation of free water in such large datasets, along with the wealth of clinical data available, including pharmacological and surgical treatment responses, quality of life measures, and survival rates, could be crucial in assessing neoplastic infiltration and edema, potentially providing new radiomic features for tumors. This highlights the need for free water estimation that is accurate in the presence of peritumoral edema using clinical acquisitions.

The fitting of a bi-compartment model to single-shell data is a mathematically ill-posed problem [8] with infinitely many solutions. The earliest attempts of fitting a bi-compartment model relied on multiple shells [9, 10]. Pasternak *et al.* [2] extended the model fit to single-shell acquisitions by positing that the problem could be addressed by an appropriate initialization of the model parameters and by a spatial regularizer that stabilizes the fit. This free water estimation allowed for a better reconstruction of healthy fornix tracts and enabled fiber tracking through edema. The application of this model has led to significant findings in various applications, including depression [11], Parkinson’s disease [12–14] and schizophrenia [15]. The initialization, however, is affected by scanner inhomogeneity [16] and it underestimates the free water in healthy tissue while producing physiologically implausible diffusion indices in the peritumoral region [17]. It has also not been extensively validated in simulated and brain tumor data.

The overarching goal of this paper is to present a paradigm for free water elimination in peritumoral tissue to obtain diffusion indices in the presence of edema, subsequently leading to improved tractography, using clinically acquired single-shell data. This is achieved by fitting a bi-compartment model, extending the work in [2] with a novel interpolated initialization designed to work optimally in edematous and healthy regions. The two compartments describe the underlying tissue and edema. This free water estimation paradigm will subsequently be referred to as FERNET (Freewater EstimatoR using iNtErpolated iniTialization). FERNET will be tested on simulated DTI data with varying free water fractions, anisotropy levels, and underlying diffusivities, and in data of healthy participants and tumor patients.

## 2. METHODS

### 2.1. Overview

We describe a novel initialization approach, aiming to improve free-water estimation from clinically acquired single-shell data, in both healthy tissue as well as in peritumoral regions. We present comprehensive experiments on both simulated and real data, demonstrating the performance of our method compared to existing ones in both healthy and peritumoral tissue.

### 2.2. Freewater EstimatoR using iNtErpolated iniTialization (FERNET)

Following the bi-tensor approach suggested previously [2, 10], we model the diffusion signal in each voxel with two compartments: a tensor compartment representing the underlying tissue and an isotropic compartment with a fixed diffusivity equal to the diffusivity of free water in biological tissues. This is mathematically modeled as

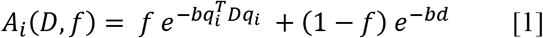

where the first term models the tissue compartment and the second term represents the free water compartment. *A*_*i*_ is the signal attenuation of the diffusion weighted image acquired along the *i*^*th*^ gradient direction, *f* is the tissue volume fraction, *b* is the magnitude of diffusion weighting, *q*_*i*_ is *i*^*th*^ gradient direction, *D* represents the diffusion tensor used for modeling the tissue compartment, and *d* is the diffusivity in the isotropic compartment, which is fixed at 3.0 × 10^−3^mm^2^/s. Fitting this model using a single-shell dMRI acquisition is a problem with infinitely many solutions [2]. Finding a solution to such a problem requires a combination of a good initialization and optimization. We use a novel initialization designed to address both healthy and pathological tissue followed by gradient descent to fit the model. Specifically, we propose the following initialization of the tissue volume fraction:

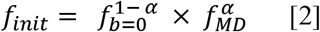

*f*_*init*_ is a log-linear interpolation between two initialization strategies, one denoted as *f*_*b*=0_ and the other as *f*_*MD*_.

The first strategy, *f*_*b*=0_, is similar to that proposed in [2], based on scaling the mean unweighted image (*S*_0_) with respect to representative unweighted signals of WM (*S*_*t*_) and CSF voxels (*S*_*w*_):

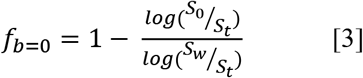

The resulting map *f*_*b*=0_ is further constrained by estimates of the minimum and maximum tissue fraction corresponding to minimum and maximum expected eigenvalues (i.e., diffusivities) of a tensor modeling brain tissue, given by

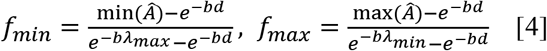

where *λ*_*min*_ and *λ*_*max*_ are set to 0.1×10^−3^ mm^2^/s and 2.5×10^−3^ mm^2^/s, respectively, and *Â*, is the vector of signal attenuation, *i.e.* the values of the diffusion weighted images divided by *S*_0_.

There are two differences in our proposed estimation of *f*_*b*=0_ from the previously reported method for free water elimination [2]. First, we define *S*_*t*_ as the 5^th^ percentile of the unweighted signal in a region of WM, and we define *S*_*w*_ as the 95^th^ percentile of unweighted signal in a region of CSF. Previously, these values were defined as the mean unweighted signal within the two regions. Second, initial values outside the range [*f*_*min*_, *f*_*max*_] are set to the nearest of the two values. Previously, the estimated value was replaced with the value 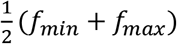 in this situation. The significance of these changes in the initialization is demonstrated in the experiments described in section 2.4.5.

The second strategy, *f*_*MD*_, is given by

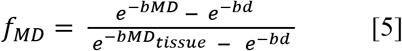

where MD is the mean diffusivity from the standard tensor fit in a voxel of interest, and *MD*_*tissue*_ is a fixed value (0.60×10^−3^ mm^2^/s [18, 19]) representing the expected MD of a WM voxel that is not impacted by partial voluming or pathology.

Finally, the value of *α* in Equation 2 is set following Equation 3, but constrained to the range [0,1] rather than the range [*f*_*min*_, *f*_*max*_]. This results in an interpolation of the two strategies which depends on *S*_0_ in the voxel of interest. For example, in a WM voxel with normal appearing T2, where the *S*_0_ is close to the value *S*_*t*_, the value of *α* is nearly 1 and the result of this interpolation is closer to that using *f*_*MD*_. In regions that appear like CSF in the T2 contrast, the value of *α* is nearly 0 and the result is closer to *f*_*b*=0_. In voxels with intermediate T2 intensity, which includes vasogenic edema, the peritumoral region, and voxels with partial volume of CSF, this interpolation is closer to the geometric mean of the two approaches. In this way, our initialization is designed to smoothly modulate its behavior based on the signal properties of the voxels without relying on any segmentation beyond the selection of representative CSF and WM regions. Because the proposed method depends on the *S*_0_ in each voxel, any bias in that image resulting from inhomogeneous magnetic field in the scanner will likely lead to a biased *f*_*init*_ (as well as a biased *f*_*b*=0_). Therefore, we propose the use of bias-field-corrected maps of *S*_0_ for free water elimination problems. The significance of the bias field correction will be demonstrated in section 2.4.5.

In summary, our interpolated initialization method (FERNET) differs from the approach in [2], which we subsequently call “b0 initialization”, in three key aspects:

- A novel initialization strategy that aims to improve the estimation of the free water compartment in both healthy and pathological tissue.
- Bias field correction of *S*_0_ before initialization.
- The initial fraction values outside of the plausible range are replaced with the nearest plausible value rather than the mean before optimization.

### 2.3. Experiments on simulated data

#### Creation of simulated data

To address the lack of ground truth, especially in edematous and infiltrated regions, we simulated data with varying ground truth mean diffusivities, anisotropies, and free water volume fractions. These simulated datasets follow a bi-compartment model where one of the compartments represents tissue and the other is isotropic with a fixed diffusivity (3.0 × 10^−3^ mm^2^/s). The simulation of the weighted images was obtained using the bi-tensor model available in Dipy’s multi-tensor simulator [20]. For simulating an unweighted (b0) image, S_0_, we calculated the transverse magnetization of a spin-echo experiment as a linear sum of the contribution of each compartment as proposed in *Phantomαs* [21]:

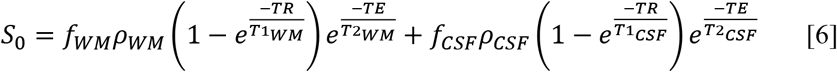

where *f*_*WM*_ is the volume fraction of white matter, *f*_*CSF*_ is the volume fraction of CSF, *ρ* is the proton density, and TR/TE are repetition and echo times. The first term of Equation [6] represents the signal from WM, while the second term represents the signal from CSF. Although values of *ρ*, T1 and T2 of the two tissue types are not known *a priori*, they do not change when volume fraction is varied. Thus, we simulate *b*_0_ for each voxel as:

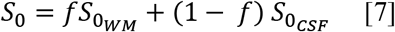

where *f* is the desired volume fraction of tissue, and 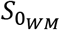 and 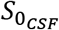 are values taken as reference from selected voxels in human data. The diffusion signal, *S*_*i*_, for a given voxel is simulated as:

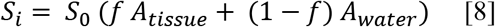

where *A*_*tissue*_ and *A*_*water*_ represent the attenuation of tissue and water, respectively. We modeled tissue with varying degrees of free water contamination and ground truth diffusion tensor indices. FA and MD values of three selected voxels were used as ground truth FA and MD values for the simulation, and their corresponding b0 values were used to simulate varying levels of volume fraction.

We simulated three clinically relevant scenarios: a) vasogenic edema, where the underlying tissue structures were preserved, i.e., a simulated “normal” white matter tissue voxel contaminated by water (MD=0.60 × 10^−3^ mm^2^/s, FA=0.6); b) vasogenic edema, where the underlying white matter tissue had higher diffusivity (MD=0.86 × 10^−3^ mm^2^/s, FA=0.7); and c) cytotoxic edema, where the underlying tissue had lower diffusivity (MD=0.40×10^−3^ mm^2^/s, FA=0.2). In each scenario, we added a free water compartment whose volume fraction varied from 0 to 1, and ten levels of Rician noise (signal to noise ratio (SNR) of 10 to 90, as well as noise-free), with 100 simulations per noise level. In addition, for every simulated tensor, we rotate the tensor directions randomly 100 times. Thus, there were 1,000,000 simulated tensors and free water compartments for each of the three scenarios. For more details on the parameters used for simulations, see Table 1.

**Table 1.** Parameters selected for simulating data to validate the proposed method

|  | Range of values | Number of simulations |
| --- | --- | --- |
| MD values | (0.4 - 0.86) x 10<sup>-3</sup> mm<sup>2</sup>/s | 3 |
| FA values | 0.2 - 0.7 | 3 |
| Free water volume fractions | 0 - 0.9 | 10 |
| Rotation of the tensor directions | Random | 100 |
| Rician noise | 10 - 90 (in steps of 10),<br>noise-free | 1000 |
| Total |  | 3,000,000 |

#### Free water estimation on simulated data

We assess our proposed method using ground truth simulated free water values as a reference point, by comparing the free water estimation following FERNET with the estimation following the b0 initialization. For all simulations, we calculated the error of free water estimation as the difference between ground-truth free water and estimated free water. In each scenario, the error in free water volume fraction was calculated when the ground truth free water was changed from 0 to 0.9. Additionally, the effect of SNR was investigated at a free water volume fraction of 0.4.

### 2.4. Experiments on human data

#### 2.4.1 Datasets

##### Dataset 1

32 participants were selected from an ongoing tumor study, including 23 healthy controls, and 9 patients with a diagnosis of glioblastoma. All participants underwent a multi-shell diffusion acquisition with b-values of 300 (15 directions), 800 (30 directions), and 2000 s/mm^2^ (64 directions) and 9 unweighted volumes. The data were acquired using a Siemens 3T TIM Trio scanner with a 32-channel head coil and echo planar (EP) sequence with TR=5216 ms, TE=100 ms, at a spatial resolution of 2 × 2 × 2 mm. T1, T2, FLAIR, and T1 contrast-enhanced (T1CE) scans were also acquired for the 9 brain tumor patients.

##### Dataset 2

This dataset comprised 143 brain tumor patients (89 glioblastoma/54 metastasis), putatively representing a range of levels of tumor cell infiltration and extent of edema. The age range of this data was 19 to 87 (77 females). Single-shell diffusion data were acquired on Siemens 3T Verio scanner with TR=5000ms and TE=5000/86ms, 3 unweighted volumes, and 30 diffusion weighted volumes at a b-value of 1000 s/mm^2^. The acquired spatial resolution was 1.72 × 1.72 × 3 mm. Additionally, T1, T2, FLAIR, and T1CE were acquired.

##### Dataset 3

This dataset comprised 216 healthy controls from the Philadelphia Neurodevelopmental Cohort (PNC) [22]. The age range of these selected subjects was 18 to 23 years. There were 118 females and 98 males. The data were acquired on a 3T Siemens TIM Trio scanner, using a 32-channel head coil and a twice-refocused spin-echo (TRSE) single-shot EPI sequence with TR=8100 ms and TE=82 ms, at a b-value of 1000 s/mm^2^ for a total 64 weighted diffusion images and seven unweighted images. The acquired spatial resolution was 1.875 × 1.875 × 2 mm.

##### Data preprocessing

All datasets were investigated visually for quality assurance (QA). The datasets were denoised [23] and corrected for eddy current-induced distortions [24]. Voxels were resampled to 2mm isotropic resolution. The b0 images were extracted and skull-stripped using FSL’s BET tool [25]. Bias correction of the b0 images was performed with N4BiasFieldCorrection from the Advanced Normalization Tools (ANTs) package. A DTI model was fit within the brain mask using an unweighted linear least-squares method. FA and MD maps of healthy controls in *Dataset 1* and *Dataset 2* were non-linearly registered to the JHU-MNI-ss (“Eve”) atlas [26] using the SyN registration algorithm from the ANTs package. Automatic tumor and peritumoral region segmentation was obtained in the patients from *Dataset 1* and *Dataset 2* using GLISTR [27], after co-registering the T1, T2, FLAIR and T1CE images using FSL’s flirt tool.

#### 2.4.2. Comparison of single-shell and multi-shell free water estimations

Since ground truth values of the free water volume fraction in human data are not available, we used the values estimated through a multi-shell estimation [28], utilizing all the shells available in *Dataset 1*, as a reference standard. We further extracted the b=800 s/mm^2^ shell to obtain single-shell data on which the two initialization approaches for free water estimation were applied. Two analyses were performed. First, the single-shell methods were compared to the reference standard by calculating voxel-wise correlation coefficient and mean square error, using the 23 healthy controls which were co-registered to the same atlas. The second analysis was performed in the peritumoral regions of 9 tumor patients to compare the estimated free water of the two single-shell approaches against the reference standard.

#### 2.4.3. Free water estimation in brain tumor data

We selected two tumor patients from *Datatset 2*: Patient 1 with a metastatic tumor and Patient 2 with glioblastoma. These represent case studies for investigating the results of the two approaches. We visually compared the free water and corrected FA of the peritumoral region to healthy tissue in the contralateral hemisphere.

#### 2.4.4 The effect of regularization on free water estimation

In order to demonstrate that our proposed initialization alleviates the dependence on regularization, for Patient 2 discussed in 2.4.3, we compared the histogram of the free water values in the whole brain as well as in the peritumoral region with and without the use of regularization for the two initialization approaches.

#### 2.4.5. The effect of bias field correction and constraints on free water estimation

As shown in Equation 3, the initialization depends on representative b0 signals in both tissue and free water compartments as well as the b0 signal itself. We applied a bias field correction [29] on the b0 image, and compared results of free water estimation with and without the application of bias correction. Analysis was performed using co-registered data from *Dataset 3*. At every voxel, the percentage of controls with artifactual initial estimates of the free water corrected tensor, defined as MD < 0.40×10^−3^ mm^2^/s [30], was calculated. In addition, we investigated the effect of constraining the initialized free water values to the extremes of the range [*f*_*min*_, *f*_*max*_], rather than the value 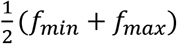, as described in section 2.2.

#### 2.4.5. Tractography

Tractography was performed on *Dataset 2* (143 brain tumor patients) using Diffusion Toolkit [31], on tensor images calculated with and without FERNET free water elimination. Tensors calculated without free water elimination are referred to as the “standard” tensor fit. The second order Runge-Kutta algorithm was used for tracking, with an angle threshold of 45°, a step size of 1 mm, and an FA threshold of 0.2. Five bundles of interest (corticospinal tract, inferior longitudinal, inferior fronto-occipital, uncinate and arcuate fasciculi) in each hemisphere were extracted from each tractogram, using the RecoBundles algorithm [32], with a pruning parameter of 7 mm. Finally, the “edema coverage”, defined as the percentage of voxels in the peritumoral edema region containing one or more streamlines belonging to any of the ten bundles of interest, was computed for every patient with and without free water elimination. A percentage difference was calculated relative to the edema coverage of the tractography without free water elimination.

## 3. RESULTS

### 3.1. Free water estimation on simulated data

Fig. 1 presents the results of free water estimation on three simulated scenarios: cytotoxic edema with underlying pathological tissue (Fig.1 A), vasogenic edema with underlying healthy tissue (Fig.1 B), and vasogenic edema (Fig.1 C). Our findings show that in the presence of simulated edema (FW>0.3), FERNET improved the estimation of the free water across the three scenarios (Fig. 1). The FERNET estimation had less variance than the b0 initialization, as shown by the smaller spread in the violin plots in Fig.1 A (FW=0.4, FW=0.5), Fig.1 B (FW=0.6, FW=0.7), and Fig.1 C (FW=0.6, FW=0.7). Moreover, in the case of simulated healthy tissue (Fig.1 B, where FW<0.3), we observed improved estimation of the free water compartment. There was no effect of SNR on the mean free water estimation in each of the three scenarios (Fig.1, A-C, 2^nd^ row). As expected, the variance in free water estimation decreased with increasing SNR.

**Figure 1:**
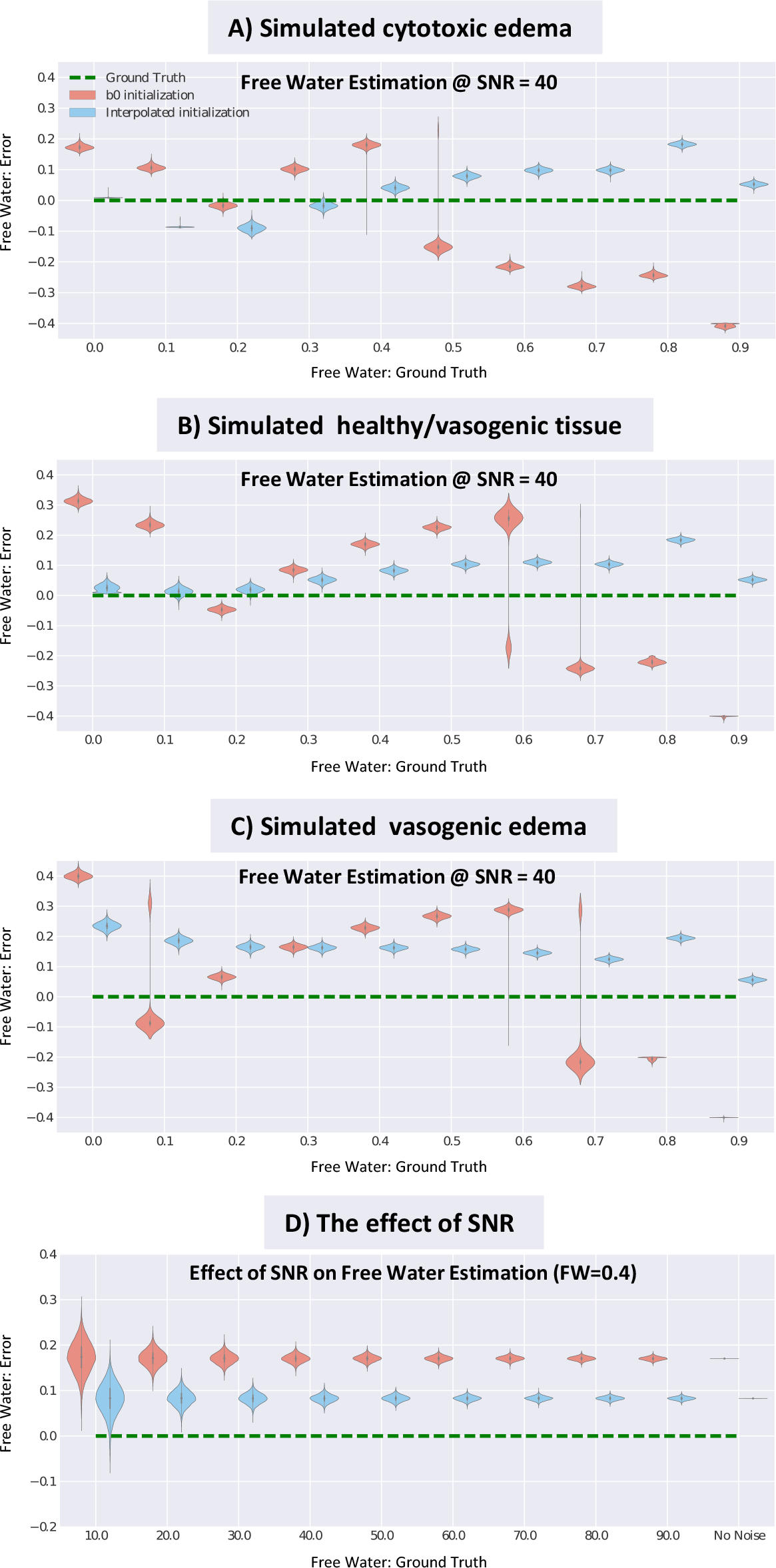
Free water estimation on simulated data. The error in free water estimation using simulated data with varying ground truth volume fraction is presented using violin plots. A) The simulated MD of 0.4×10^−3^ mm^2^/s represents cytotoxic edema. Two models of vasogenic edema are simulated: B) a simulated MD of 0.6×10^−3^ mm^2^/s approximately equivalent to that of a healthy WM voxel, and C) a simulated MD of 0.86×10^−3^ mm^2^/s higher than that of healthy WM. All experiments in the first rows have an SNR of 40, with free water varying from 0 to 0.9. The y-axis represents the difference between the free water estimation and the corresponding ground truth of free water (x-axis). Green dotted lines in each plot represent the zero line, or perfect free water estimation. Violin plots of the two methods are staggered for clarity. Results show that FERNET initialization is more accurate in estimating free water than the b0 initialization where FW>0.3. D) shows the effect of SNR on the error in free water, where SNR varies from 10 to 90, followed by a simulation with no noise. Although SNR has an effect on the variance of error in free water, the effect of SNR on the mean error in free water is negligible.

### 3.2. Free water estimation on human data

Fig. 2 shows the results of comparing the free water estimation using FERNET and the b0 initialization with the free water estimation obtained from the multi-shell method used as a reference standard in the absence of ground truth. Voxel-wise maps of correlation coefficient and mean square error (MSE) of the free water fraction between single-shell methods and the multi-shell method (Fig. 2 A for the b0 initialization, Fig. 2 B for the interpolated initialization of FERNET), showed a weaker correlation between the free water maps, especially in WM regions, and a relatively higher MSE in the b0 initialization when compared to FERNET. These correlation findings were consistent with Fig. 2 C, where the free water values in WM of 23 healthy controls using FERNET were more aligned with the identity line when compared to the multi-shell reference standard. The correlation between b0 initialization and multi-shell estimation was 0.39, and the correlation between FERNET and multi-shell estimation was 0.76. The correlation of free water values in the peritumoral regions of 9 patients (Fig. 2 D) between the b0 initialization and the reference standard was 0.45, and between FERNET and the reference standard was 0.71.

**Figure 2:**
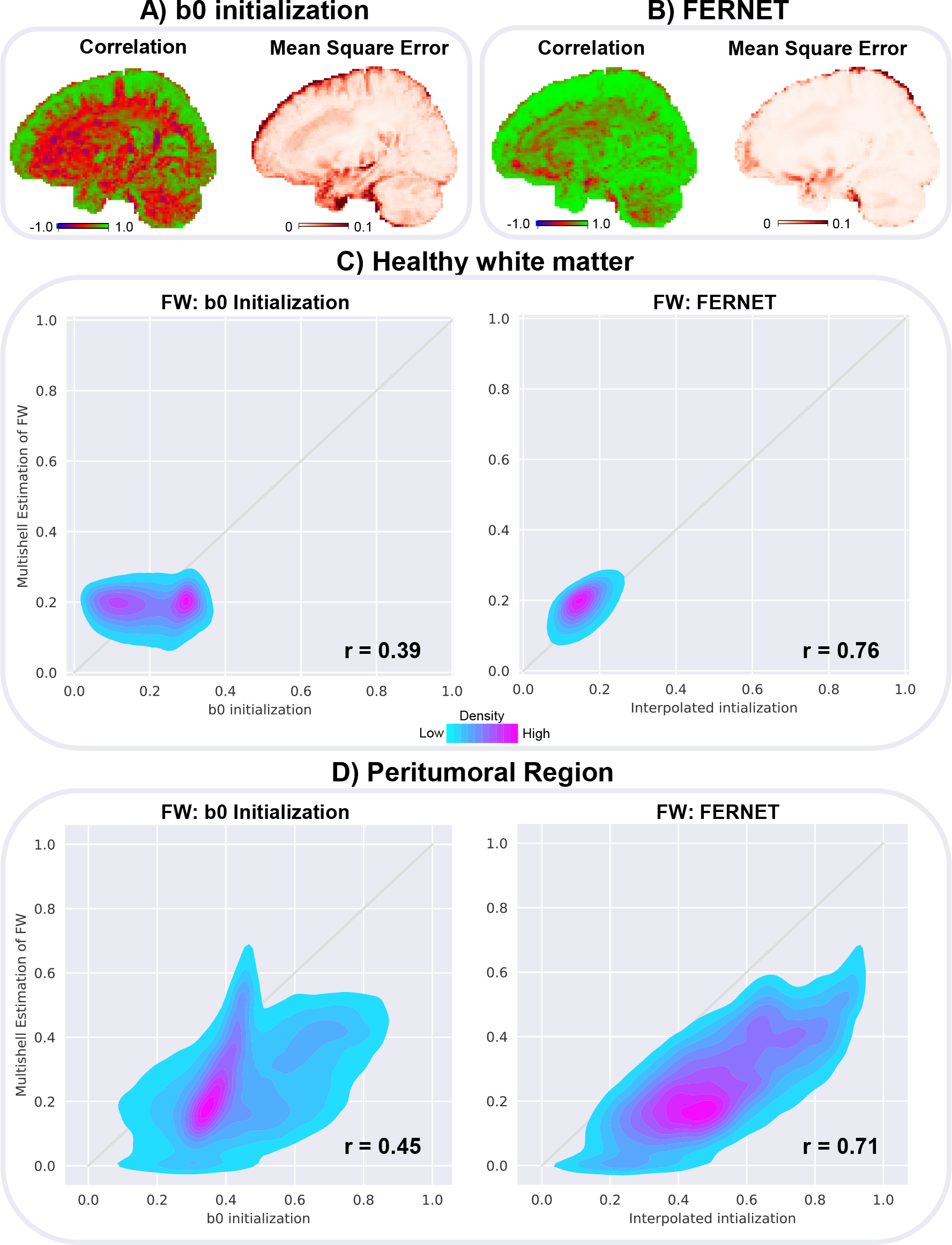
Free water estimation in human data compared to the multishell reference standard of Hoy et. al [28]. Correlation coefficient and mean square error (MSE) are measured at every voxel for 23 healthy controls to assess the contrast and difference between the free water (FW) map derived from A) b0 initialization and B) interpolated initialization FERNET compared to the FW map derived from multi-shell. C) shows the scatterplots of free water fraction values within the WM of 23 controls, demonstrating that the proposed interpolated initialization is more aligned with the reference standard in WM. D) shows the scatterplots of free water fraction in the peritumoral regions of 9 brain tumor patients, demonstrating that neither of the single-shell methods obtained an estimation in the peritumoral region that is perfectly consistent with reference standard. However, correlations are higher with FERNET (r=0.71) than with b0 initialization (r=0.45).

### 3.3. Free water elimination in brain tumor data

Fig. 3 shows the free water volume fraction and the corrected FA maps using the two free water estimation methods on two brain tumor patients: Patient 1 with brain metastasis and Patient 2 with glioblastoma. FERNET free water maps had spatially smoother contrast than those from the b0 initialization, and FA values in the WM ipsilateral to the tumor matched the FA of the contralateral WM better, especially in Patient 1.

**Figure 3:**
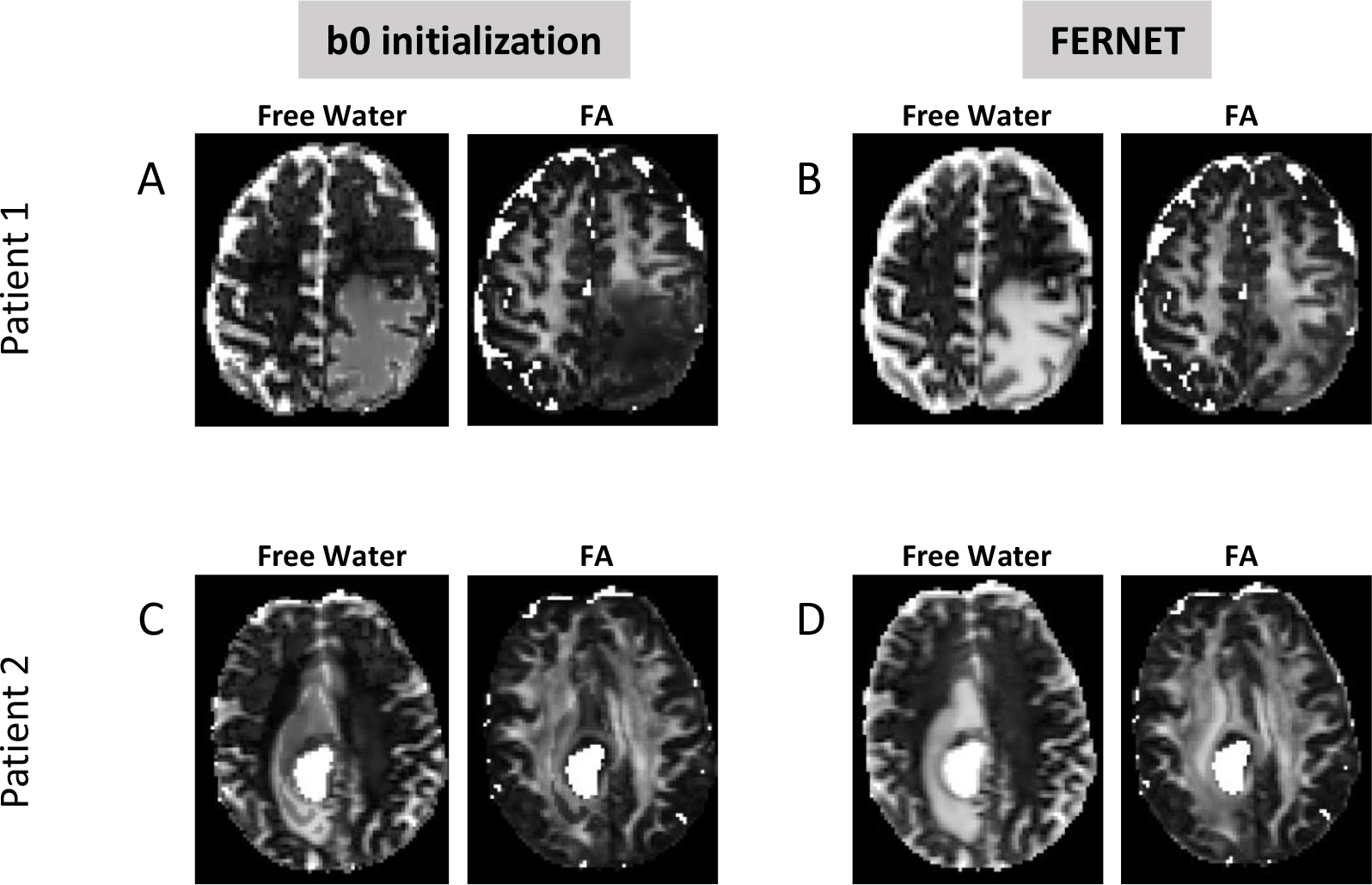
Comparison of free water estimation and corrected FA maps between the two initialization methods in two brain tumor patients. Patient 1 is a metastatic brain tumor patient and Patient 2 is a glioblastoma patient. Corrected FA maps obtained with FERNET (B, D) show better agreement between the peritumoral region and the contralateral WM compared to b0 initialization (A, C). Free water volume fraction maps obtained with FERNET are spatially smoother in the peritumoral region.

### 3.4. The effect of regularization on free water estimation

Fig. 4 shows the behavior of the free water estimation approaches to regularization in two brain tumor patients. The effect of regularization is assessed via the difference maps of the free water volume fraction obtained with and without regularization, in the peritumoral area and the whole brain. The histograms of the free water estimation in the whole brain and in the peritumoral region show a closely matched free water estimation for regularized and non-regularized approaches when using FERNET.

**Figure 4:**
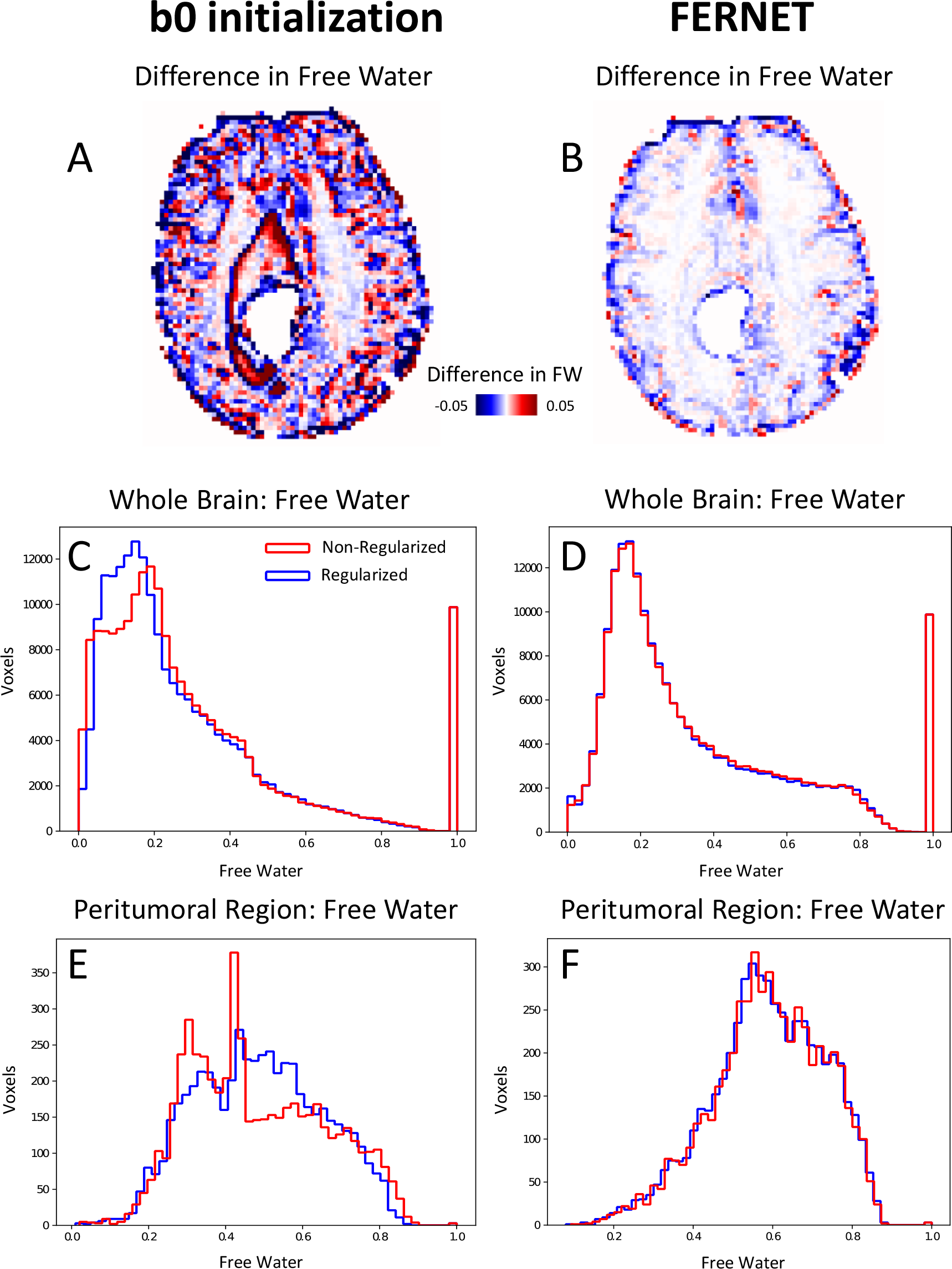
Effect of regularization in the optimization phase after FERNET and b0 initialization in a tumor patient. A and B show the difference images of free water volume fraction between regularized and non-regularized fits of the b0 initialization method and FERNET, respectively. Histograms of free water volume fraction in the whole brain with and without regularization using b0 initialization differ greatly (C), while the histograms of free water using FERNET closely match (D). Histograms in E and F demonstrate a similar effect when limited to the peritumoral region.

### 3.5. Effect of bias field correction and constraints on initial free water

Fig. 5 shows the results of investigating the use of bias field correction in free water elimination. It shows the percentage of participants that yielded an artifactual fit (MD<0.40×10^−3^ mm^2^/s) in the initial corrected tensor (before the optimization phase) at each voxel in the brain, with (Fig. 5 A) and without (Fig. 5 B) bias field correction. Without bias field correction, the corrected tissue tensor is initialized with indices that are physiologically implausible. Such diffusivity values are estimated in a large percentage of controls in the peripheral regions of the cortex and cerebellum, areas which are most impacted by bias. Finally, Fig.5 C shows that constraining the initial free water to the range [*f*_*min*_, *f*_*max*_] rather than replacing with the mean value (see section 2.2) leads to a more plausible contrast in the free water volume fraction map, more similar to the FLAIR contrast.

**Figure 5:**
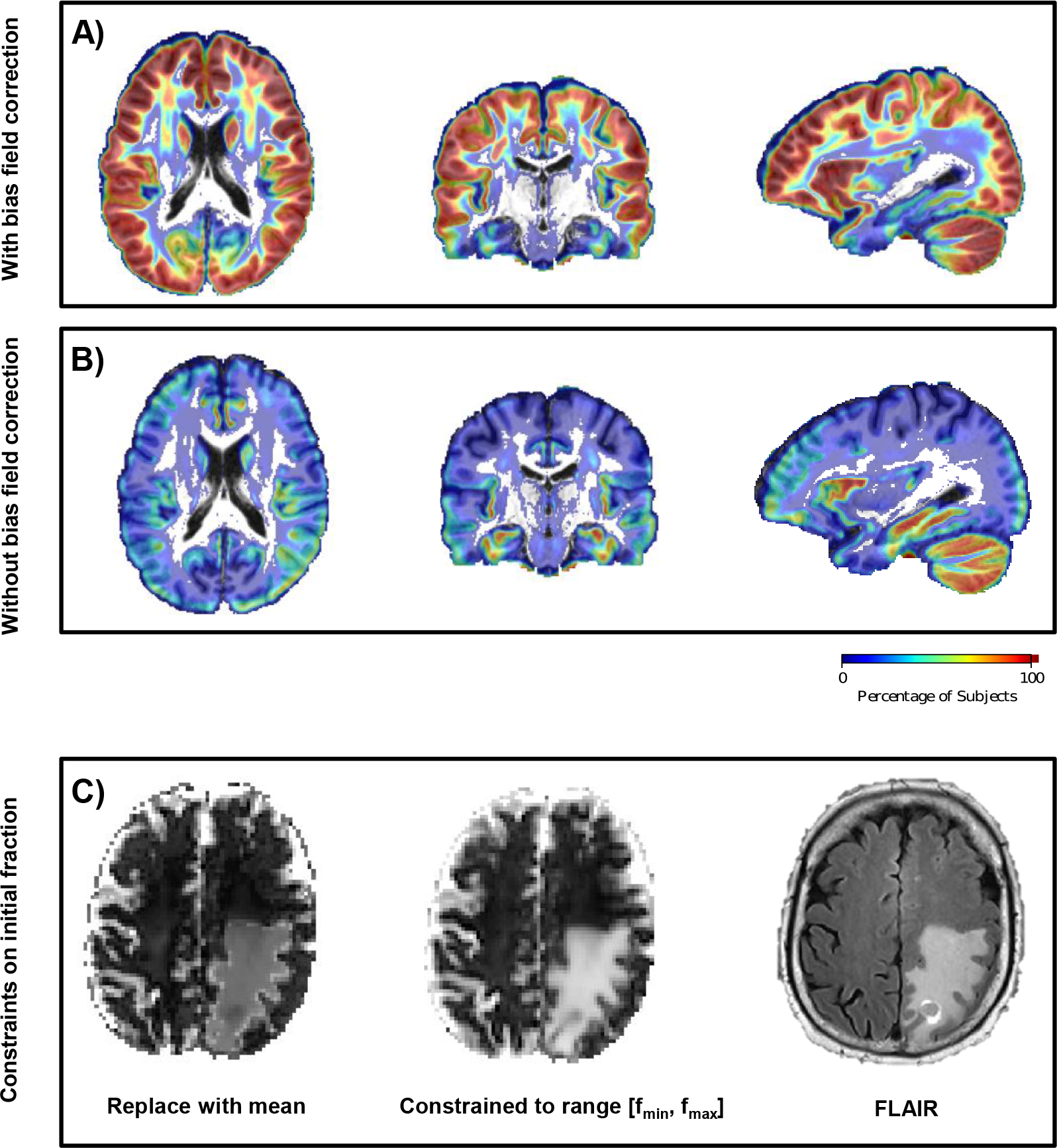
Effect of bias field correction and heuristics on corrected diffusion indices and free water volume fraction. Evaluation of FERNET and b0 initialization on a large dataset (*Dataset 3*) of healthy controls showed that A) data without bias field correction produced more physiologically implausible voxels (defined as corrected MD < 0.4×10^−3^ mm^2^/s) using FERNET than data with bias field correction (B), especially in GM regions and in the cerebellum, which are the areas impacted most by bias. The colorbar represents the percentage of controls with physiologically implausible fits. C) Constraining initial free water estimation to the boundaries of feasible free water values instead of the mean value (see Equation 4) yields a free water map in the peritumoral region that is more plausible and more aligned with the FLAIR contrast.

### 3.6. Tractography

Fig. 6 shows the effect on tractography as a result of free water elimination in a large dataset of 143 brain tumor patients (*Dataset 2).* Fig. 6A is an example case, showing a comparison of the arcuate fasciculus between standard tensor-based tracking and FERNET-based tracking. In extending this tractography to all the patients, we found that the tractography based on FERNET covered more of the peritumoral region than tractography using the standard tensor fit. This is depicted in Fig. 6 B by a histogram of percent difference in coverage of the edematous region by tracts. In a majority of patients, the percent difference in edema coverage was positive, indicating that tractography on free-water-corrected tensor maps traveled through a larger portion of the peritumoral region.

**Figure 6:**
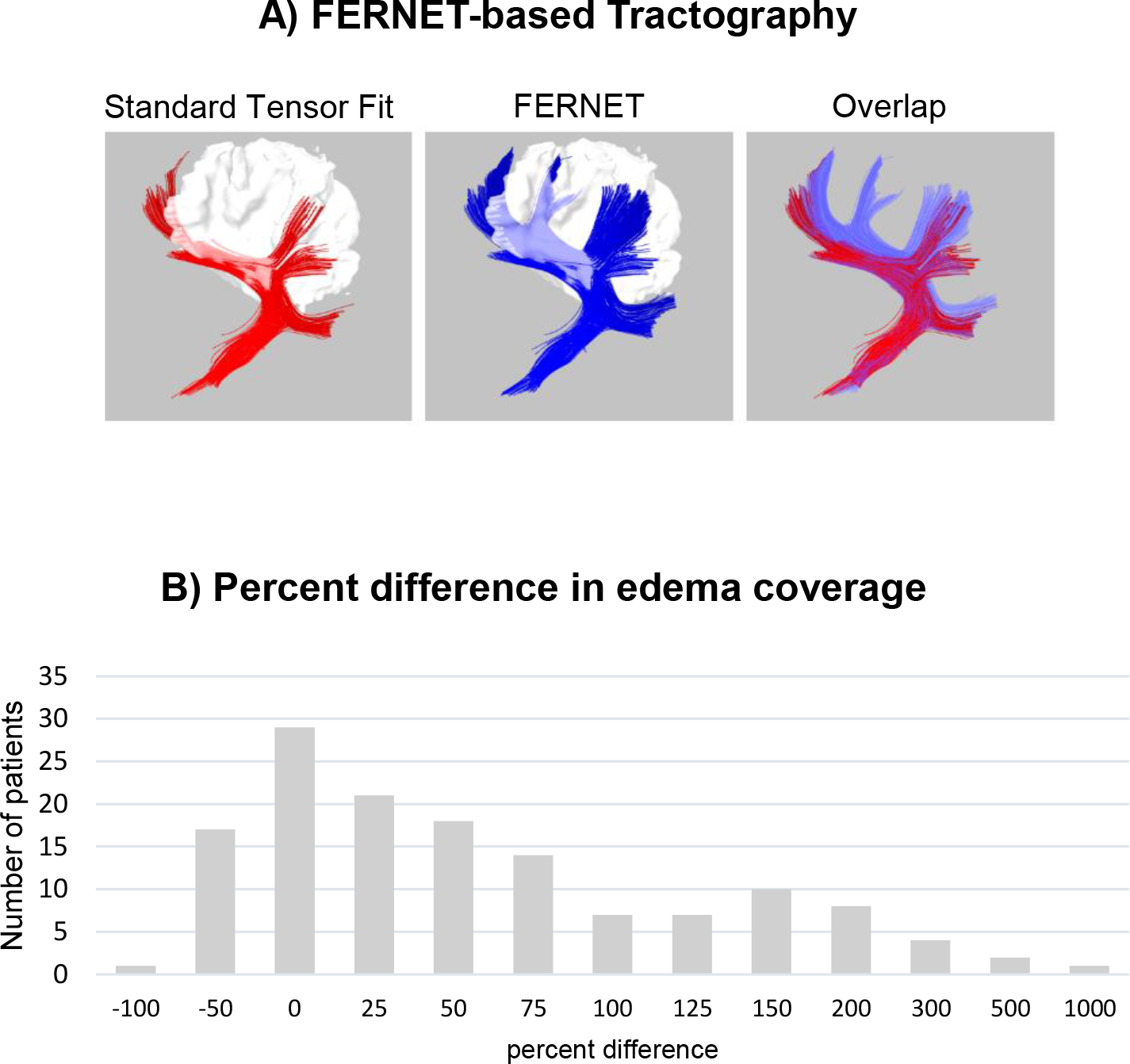
Fiber tractography with and without free water elimination using FERNET. A) Tractography of the arcuate fasciculus in one example brain tumor patient demonstrates the improvement in tracking inside the peritumoral region using free water elimination. Using standard tensor fit based tracking, streamlines appear to stop prematurely when they reach voxels affected by the accumulation of free water in the peritumoral region. The overlap of both approaches is shown, highlighting the improvement observed with FERNET. B) Tractography of ten major tracts in 143 tumor patients showed an increase in the volume of the peritumoral region traversed by streamlines, as shown in a histogram of percent difference (relative to standard tensor-based tractography) in edema coverage. The findings demonstrate that most patients exhibit an increase in edema coverage when free water elimination is employed.

## 4. DISCUSSION

We have introduced a new initialization strategy, FERNET, for the free water elimination problem in single-shell data, aimed at improving the characterization of peritumoral edema using clinically feasible diffusion MRI. The efficacy and wide applicability of our method in estimating free water fraction has been demonstrated comprehensively both on simulated data representing vasogenic and cytotoxic edema, as well as datasets of healthy controls and tumor patients of different tumor types. We have shown that our strategy improves the accuracy of free water elimination for peritumoral regions.

Most free water elimination methods to date have been designed for healthy tissue. Developing a method for a pathology, like tumors, is very challenging due to the absence of ground truth. We aimed to address this issue by generating simulated data representing vasogenic and cytotoxic edema using different ground truth diffusivities, anisotropies, and free water volume fractions, and tested our proposed interpolated initialization against the existing b0 initialization method. Fig. 1 highlights the significant impact of the initialization strategy on the free water estimation using three simulated clinically relevant scenarios. The findings show that the proposed initialization strategy yields improved free water estimation, especially in voxels affected by simulated “edema”. Improvement in free water estimation was also observed in the “healthy” tissue. Also, we showed (Fig.1 B and C at simulated FW=0.6 and FW=0.7) that the interpolation approach provides a robust estimation that is less susceptible to noise. Although, as expected, the free water estimation became increasingly robust as SNR increased, the effect of SNR on the mean of free water estimation was negligible (Fig.1 D).

In order to translate the findings in Fig.1 to human data, we used the free water estimated by Hoy et al. [28] as a reference standard for free water in human data, as no ground truth exists, especially when tissue is contaminated with vasogenic and cytotoxic edema. The estimation in this method relies on multi-shell data and claims to have a closed form solution to a problem that is ill-posed when attempted using single-shell data. The voxel-wise correlation and MSE maps in Fig. 2 A and B show that FERNET based free water maps, although using a single shell, are more similar in contrast to the reference standard, and they correlate better with the free water estimation of this advanced method compared to b0 initialization. This increased agreement of FERNET, as seen in Fig. 2 C and D, suggests the superiority of FERNET initialization in the peritumoral region and also in healthy tissue.

There were several methodological choices associated with free water elimination using FERNET. As free water elimination using single-shell DTI is an ill-posed problem, previous approaches (Pasternak et al. 2009) have emphasized the importance of regularization. Fig. 4 A shows that FERNET initialization is less sensitive to regularization, as shown in difference images between regularized and non-regularized optimization following FERNET initialization and b0 initialization. This implies that FERNET yields a solution that varies smoothly across neighboring voxels. On the other hand, the estimation using b0 initialization appears to be more influenced by regularization, with most impacted regions in healthy-appearing GM and the boundaries of the peritumoral region. This effect is further illustrated by the histograms of free water fraction with and without regularization in Fig. 4 C-F.

The efficacy of choices we made in the implementation details of our initialization was further demonstrated in Fig. 5. In Fig.5 A and B, we demonstrated that applying bias field correction to the data reduced the prevalence of implausible diffusivity values in a large set of healthy controls, which were defined as having MD less than 0.4×10^−3^ mm^2^/s, a value that is well below most estimates of the MD of healthy brain tissue. This finding suggests that, in the presence of magnetic field inhomogeneity in the scanner, the preprocessing of diffusion data may play a significant role in modeling the diffusion signal with this bi-compartment model, and it must be approached carefully. Fig. 5 C exhibits the effect of constraining our initial volume fraction estimate to the boundaries ([*f*_*min*_, *f*_*max*_]) and scaling the b0 image by the 5^th^ and 95^th^ percentile of the unweighted signals of WM and CSF, respectively. As can be seen in the figure, the FERNET initialization resulted in a contrast in the volume fraction map that was consistent with that obtained from FLAIR images. FLAIR is the clinical modality of choice to draw inferences about the peritumoral region, because in FLAIR images the inversion time is carefully chosen to null the signal from the CSF while increasing the signal intensity in edema and areas of inflammation with increased partial voluming of fluid. This suggests that the free water maps obtained with FERNET may have clinical relevance in locating and characterizing lesions and areas of inflammation such as in the peritumoral region, multiple sclerosis lesions, or inflammation caused by traumatic brain injury.

The clinical significance of FERNET is underscored by examining the free water volume fraction maps in different types of tumors that may have varying degrees of vasogenic and cytotoxic edema. The findings of Fig. 3 in the peritumoral region of two brain tumor patients, one with a metastatic tumor and one with a diagnosis of glioblastoma, showed that the interpolated initialization allowed for reconstructing FA in the region of WM ipsilateral to the tumor, despite the presence of edema. The b0 initialization based free water estimation struggled significantly, especially in the case of metastatic brain tumor (Patient 1). Thus, FERNET is generalizable to peritumoral regions with different types of characteristics.

Finally, tractography algorithms with their ability to delineate eloquent fiber tracts has important clinical implications for the surgical treatment of malignant brain tumors [33]. Although no current method solves the inherent limitations of fiber tractography, the techniques available are sufficiently specific and sensitive to make fiber tractography a valuable tool for the neurosurgeon [34, 35]. However, the standard tensor fit does not account for any free water compartment, and in the presence of significant partial voluming of extracellular fluid, indices derived from the fitted tensor represent erroneous estimations of the underlying tissue, including low FA. This may cause some tracking algorithms to stop prematurely (Fig. 6 A) in fluid-contaminated regions, leading to spuriously misshapen and/or broken tracts [36]. Although this is more pronounced in the presence of edema and inflammation related to liquefactive necrosis, it is important to note that even in the absence of pathology, cerebrospinal fluid (CSF) partial voluming affects the reconstruction of WM tracts such as the fornix, potentially resulting in misleading inferences about brain connectivity.

The improved free water estimation that our initialization provides leads to a better modeling of brain tissue. Hence, the elimination of the contamination of free water in the diffusion signal, whether it is due to partial voluming or pathology, results in better tractography than the standard single tensor model can provide. Our method of free water elimination (and its subsequent impact on tractography) was evaluated in 143 brain tumor patients (89 gliomas and 54 metastases), comparing standard tensor and tensors obtained after free water elimination using FERNET. Fig. 6 B demonstrates that the fiber tractography in peritumoral region is significantly improved with our method, as can be seen in the histogram of percent difference of edema coverage, showing increased volume covered by streamlines generated by tractography up to 1000% in some patients. Some participants exhibit a decrease of the coverage inside the peritumoral region, which may be due to the reduction of the spurious streamlines post free water elimination.

There is a growing evidence that the objective of “maximal safe resection” [37, 38] is best achieved with detailed mapping of the brain tumor and surrounding white matter tracts. This emphasizes the importance of free water elimination for tractography, paving the way for robust surgical planning, and superior tracking intraoperatively [39]. FERNET provides a method that can possibly enhance clinical outcomes, due to the improvement in tractography, which is expected to have a large impact on a broad spectrum of applications that use dMRI, including neurosurgical planning [40, 41], assessing connectivity changes in pathologies traumatic brain injury [42], stroke [43], and in pre-clinical studies such as mapping the structural connectivity of neural ganglia [44].

This study has a few limitations. Due to the absence of the ground truth values in human data, this study relies on an estimation of free water using a multi-shell acquisition as a reference standard. We trust that this is the best that can be done for healthy or peritumoral tissue, both of which cannot be biopsied to generate pathology-based ground truth. Our experiments on human data are amply supported by the large number of experiments on simulated healthy and pathological tissue. Furthermore, this bi-compartment model represents white matter with only a single tensor, causing tractography to be limited in the crossing fiber regions, as it is in standard tensor-based tractography. However, the observed improvement in tractography in the peritumoral region, even with clinically acquired data where higher order diffusion models cannot be fitted, underlines the importance of FERNET.

## 5. CONCLUSIONS & FUTURE WORK

We have designed a free water estimation paradigm based on a novel initialization approach, FERNET, that can be applied to single-shell diffusion acquisitions, readily available in the clinic. The initialization approach, which interpolates diffusivity and T2 information (b0), improves free water estimation in the tissue contaminated by edema and/or infiltrative neoplasm, as well as healthy tissue, compared to traditional approaches. This is a significant contribution because of the widespread prevalence of single-shell diffusion acquisitions in clinical studies.

FERNET initialization for free water estimation has a myriad of applications in the field of diffusion MRI, especially in pathologies that lead to edema, like tumors and traumatic brain injury. As it is designed to work with clinically feasible single-shell diffusion data, it can be used to improve tractography in retrospective studies using data that is no longer considered state of the art, as suggested by our demonstrated improvement in tractography in a cohort of brain tumor patients. A comprehensive investigation of different tractography algorithms, deterministic and probabilistic, is beyond the scope of this manuscript and will be investigated in the future. Our free water map delivers valuable information about the free water content of brain tissue in the presence of pathology. This may be used to distinguish different types of brain tumors and their genetic underpinnings based on the nature of the peritumoral tissue, paving the way for better radiomic markers of cancer.

The applicability of FERNET in clinically feasible acquisitions is expected to facilitate investigations of large tumor datasets like ABTC (http://www.abtconsortium.org), that have single shell acquisitions. The tractography tested here was based on single fiber estimation, and approaches with more complicated fiber models may assist in simultaneously resolving partial volume and complex fiber architecture in the context of peritumoral regions. Finally, an interesting application of this method can be in improved estimation of structural connectomes in the presence of edema, as the estimation of structural connections in the brain is expected to improve with a better estimation of the underlying tissue compartment after mitigating the effects of partial voluming and/or pathology. These avenues will be explored in future studies.

## 6. ACKNOWLEDGMENT

This research was supported by National Institutes of Health (NIH) grants R01NS096606 (PI: Ragini Verma) and R01MH108574 (PI: Ofer Pasternak), and research grant from Synaptive Medical 30071788 (PI: Ragini Verma).

